# Negative supercoiling facilitates human TOP3α-mediated decatenation of DNA braids via catenation junction melting

**DOI:** 10.64898/2026.09.23.753725

**Authors:** Sanjana Saravanan, Anno I. Koetje, Graeme A. King, David Rueda, Luis Aragon

**Affiliations:** DNA Motors Group, MRC Laboratory of Medical Sciences, London W12 0HS, UK; Institute of Structural and Molecular Biology, Division of Biosciences, University College London, London WC1E 6BT, United Kingdom; Single Molecule Imaging Group, MRC Laboratory of Medical Sciences, London W12 0HS, UK; Department of Infectious Disease, Faculty of Medicine, Imperial College London, London W12 0HS, UK

## Abstract

DNA catenation between sister chromatids arises during replication and must be efficiently resolved to ensure faithful chromosome segregation. The removal of DNA catenation is generally attributed to the type II topoisomerase TOP2α, which catalyses double-stranded DNA passage reactions and can therefore directly remove inter-DNA linkages. By contrast, the type IA topoisomerase TOP3α catalyses strand passage through transient breaks in single-stranded DNA. Here, using single-molecule optical tweezers and fluorescence microscopy, we show that negative supercoiling promotes local melting of duplex DNA at DNA-DNA crossings within braided molecules, generating single-stranded DNA that can be engaged by TOP3α. These supercoiled substrates can be efficiently resolved by TOP3α-RMI1-RMI2 (TRR), whereas relaxed DNA is resistant to TRR decatenation. Moreover, we find that RMI2 and the ssDNA binding Replication Protein A (RPA) stimulates TOP3α decatenation and demonstrate that decatenation remains viable up to 30pN of tension on the substrate. These findings reveal a mechanism by which torsional stress converts an otherwise inaccessible duplex DNA crossing into a substrate for a single-strand-specific topoisomerase. We propose that negative supercoiling can therefore direct type IA topoisomerases to topological DNA linkages by promoting local duplex melting, providing a physical mechanism that couples DNA supercoiling to the resolution of DNA entanglements.

## Introduction

DNA replication presents a topological challenge to chromosome inheritance. As the replication machinery progresses along DNA, the intertwining of the parental duplex is redistributed into torsional stress ahead of replication forks and topological linkages between newly replicated sister DNA molecules behind them ^1^. These inter-sister DNA catenations must be removed before chromosome segregation, as their persistence can compromise chromosome individualisation and, ultimately, faithful genome transmission ^2,3^. In eukaryotic cells, the resolution of DNA catenation is principally attributed to type II topoisomerases, particularly TOP2α, which transiently cleave both strands of one DNA duplex and transport a second duplex through the resulting break ^4^. This double-strand passage mechanism makes TOP2α intrinsically suited to resolving inter-DNA linkages and underlies its essential role in chromosome decatenation during mitosis ^3^.

Type IA topoisomerases operate through a fundamentally different mechanism. Rather than transporting duplex DNA through a double-stranded break, these enzymes transiently cleave a single DNA strand and catalyse the passage of another strand through the resulting gate ^4,5^. Human TOP3α, a type IA topoisomerase, has important functions in maintaining genome stability ^6^ and, together with the BLM helicase, RMI1 and RMI2, forms the BTRR complex that promotes the dissolution of double Holliday junctions and other recombination intermediates ^7–9^. Yeast Top3 can also resolve hemicatenanes and related DNA structures containing single-stranded regions ^10^. These activities not only illustrate the ability of type IA topoisomerases to eliminate topological linkages between DNA molecules, but they also reveal an important mechanistic constraint: productive strand passage requires access to single-stranded DNA.

Evidence that type IA topoisomerases can act on interlinked DNA molecules has emerged from studies across several organisms. In *E. coli*, Topo III has been shown to decatenate newly replicated daughter DNA molecules ^11^, an activity that is strongly stimulated by the RecQ helicase ^12^. *In vitro* studies with the budding yeast homologues further demonstrated that the Sgs1-Top3-Rmi1 complex can promote both catenation and decatenation of duplex DNA, with replication protein A (RPA) markedly enhancing these reactions ^13^. In the human system, BLM has similarly been shown to stimulate the ability of TOP3α to relax negatively supercoiled DNA; however, in contrast to the yeast complex, BLM-TOP3α has not been shown to support efficient catenation or decatenation of duplex DNA in bulk biochemical assays ^8^ and the decatenation activity reported to date has been confined to single-stranded DNA ^14^. Thus, although the capacity of type IA topoisomerases to resolve DNA entanglements is well established in specific contexts, the physical features that render an otherwise duplex DNA linkage accessible to their single-strand passage mechanism remain poorly understood.

DNA topology itself could provide a means of overcoming this constraint. Negative supercoiling stores torsional energy in DNA and can promote structural transitions that relieve the energetic cost of underwinding, including local disruption of base pairing and formation of alternative, non-B-form DNA structures ^15–17^. The occurrence and stability of these transitions are strongly influenced by the mechanical state of the DNA, with both torque and tension determining the balance between B-form DNA, writhe and locally disrupted duplex states ^18–20^. In intertwined DNA molecules, DNA-DNA crossings impose additional geometrical and mechanical constraints that could potentially create privileged sites for local duplex destabilisation. Negative supercoiling might therefore alter not only the global topology of an intertwined DNA substrate but also the local molecular structure of the DNA crossing itself. Whether such mechanically induced structural changes can render a duplex DNA entanglement accessible to a single-strand-dependent topoisomerase has not been established.

Here, we use single-molecule optical tweezers to investigate how DNA supercoiling influences the resolution of intertwined DNA molecules by human TOP3α in complex with RMI1 and RMI2 (TRR). By independently controlling DNA braiding, supercoiling, and tension, we find that negative supercoiling promotes local melting of duplex DNA at DNA-DNA crossings at near zero forces, generating a substrate that can be efficiently engaged by TRR. Accordingly, TRR readily resolves DNA braids when one or both intertwined molecules are negatively supercoiled, whereas relaxed duplex braids are refractory to TRR-mediated decatenation. Replication Protein A (RPA) and RMI2 further stimulate this reaction, and braid resolution remains efficient under substantial mechanical tension. Our results reveal how torsional stress can convert a duplex DNA crossing into a substrate for a single-strand-dependent topoisomerase and establish a direct mechanistic link between DNA supercoiling, local duplex melting, and DNA decatenation. More broadly, they suggest that the local mechanical state of DNA can determine the enzymatic pathway available for resolving topological DNA linkages.

## Results

### Negative supercoiling promotes local duplex melting at DNA-DNA crossings

We first asked whether the physical environment of a DNA-DNA crossing could promote local duplex destabilisation, generating the single-stranded DNA (ssDNA) required for TRR-mediated strand passage.

To examine DNA structure at individual crossings, we generated braided DNA substrates using a quadruple-trap optical tweezers system with confocal fluorescence detection (Fig. 1A), as previously described ^21–23^. Two biotinylated λ-DNA molecules (48.5 kb) with open ends (48.5 kb) were each captured between a pair of streptavidin-coated beads and verified from their characteristic force-extension (FE) behaviour (Fig. 1B, unconstrained). A defined sequence of bead movements was then used to pass one DNA under and over the other once, generating a single right-handed braid between the two molecules (Fig. 1C) ^24^. Braid formation was independently confirmed by direct visualization of the DNA using SYTOX Orange staining, and by the characteristic force changes recorded on two of the optical traps as the molecules were interwound (Fig. 1D)

**Figure 1.**
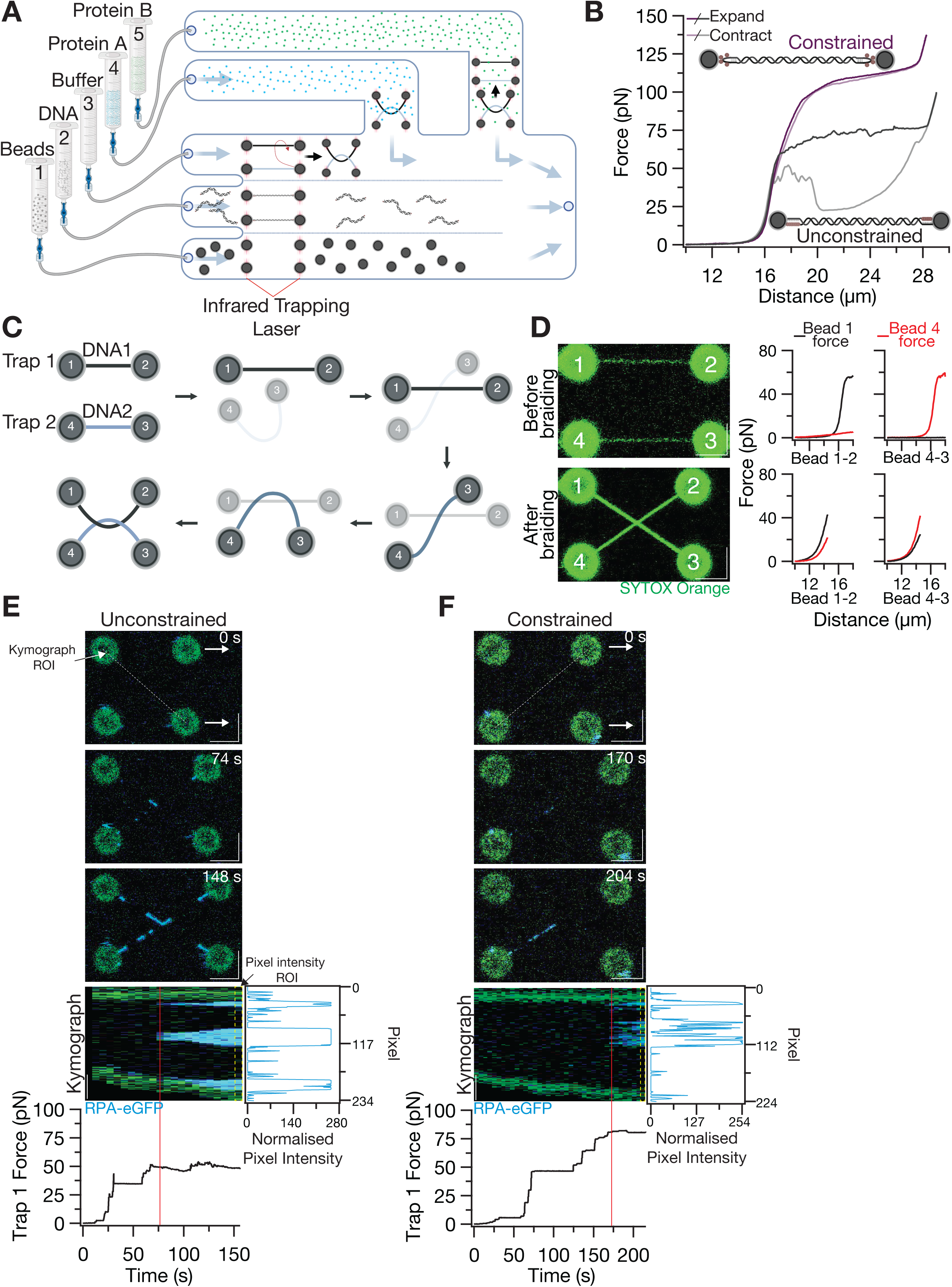
A quadruple-trap optical tweezers method for visualising dsDNA braids. **(A**) Schematic of microfluidic chip compatible with the LUMICKS C-Trap and channel contents. All experiments were conducted with the beads-, DNA-, and buffer-containing channels (channels 1-3) using 20 mM Tris-HCl pH 7.5, 50 mM NaCl buffer and the protein channels (channels 4-5) using 50 mM Tris-HCl pH 7.5, 125 mM potassium glutamate, 2.5 mM MgCl_2_, 0.5 mg/mL BSA, 1 mM DTT buffer. **(B)** Force extension (FE) comparison of a single tethered biotinylated λ-DNA that is either unconstrained (open ends) or constrained (end-capped). **(C)** Representation of the scripted steps taken to generate a single right-handed braid. Darkness of the tethered DNA and beads represents the relative position in the z-axis, with darker colours meaning the position of that trap is higher than the other trap. Trap 1 is kept on the same plane throughout the braiding process. **(D)** (left) 2-D confocal scans of DNA in the presence of 0.05 nM SYTOX Orange before braiding (held at ∼11 pN) and after braiding (held at ∼45 pN). Scale bars = 5 μm. (right) Corresponding FE curves for parallel and braided DNA. The force measured on bead 1 (Trap 1) and bead 4 (Trap 2), respectively, when either bead 2 or bead 3 is extended, before and after braiding. **(E-F)** Both DNAs in the braid are unconstrained in panel **E** and both DNAs are constrained in panel **F**. Frames at different timepoints were selected from a 2-D confocal scan taken in the presence of 5 nM RPA-eGFP. The time shown in each frame is the time to the nearest second at the start of that frame. Beads 2 and 3 were slowly extended one bead at a time in the x-axis to generate increasing forces in Trap 1 and 2. Dotted diagonal white line between beads 1 and 3 (E) and beads 4 and 2 (F) show the ROI used for the kymograph, with each pixel in the x-axis of the kymograph corresponding to a frame in the 2-D scan. Dotted yellow box designates the pixel of the kymograph used for the normalised pixel intensity plot of the blue signal. The corresponding force in Trap 1 is shown under the kymograph with the red line denoting the frame and time that the RPA signal first appears at the braid. Scale bars = 5 μm.

To investigate duplex destabilisation we monitored local strand separation using fluorescent RPA, which binds ssDNA with high affinity but interacts minimally with intact duplex DNA ^25^. RPA recruitment therefore provided a sensitive real-time readout of ssDNA exposure within individual braided substrates. We first examined braids formed from torsionally unconstrained DNA molecules, in which each duplex could rotate freely around its helical axis and show a characteristic FE profile (Fig. 1B). Using a concentration of 5 nM RPA-eGFP, RPA began to accumulate at discrete positions along the DNA under increasing force, with prominent recruitment to the DNA-DNA crossing becoming apparent only at ∼50 pN and above (Fig. 1E). Thus, sufficiently high mechanical tension can promote local duplex destabilisation at DNA crossings.

We next asked how topological constraint affected this transition. Braids were generated from torsionally constrained end-capped DNA molecules containing multiple biotin attachments at each end ^26^ (Fig. 1B, constrained). With this substrate, RPA recruitment was detected at the braid at forces above 75 pN (Fig. 1F). Therefore, while the force threshold for RPA binding at the braid was higher when both molecules were constrained, there was no significant difference in the ability of force to expose ssDNA at a DNA crossing when topological constraint is added to the DNAs in the braid.

Negative supercoiling stores torsional energy within DNA and, at forces >∼1 pN, typically favours duplex destabilisation in the form of denatured base pairing ^27^. We therefore asked whether introducing negative supercoiling could lower the force required to expose ssDNA at the crossing. Negatively supercoiled DNA molecules were generated using the Optical DNA Supercoiling (ODS) method described by King *et al*. ^28^ (Fig. 2A). This allowed us to generate braided substrates containing defined levels of negative supercoiling correlated to the difference in the extension of the DNA at 70 pN between relaxed and negatively supercoiled molecules (ΔL_70pN_) (corresponding to supercoiling densities (σ) between 0 and -0.7, with a precision of approx. 0.03) (Fig. 2B, S1A).

**Figure 2.**
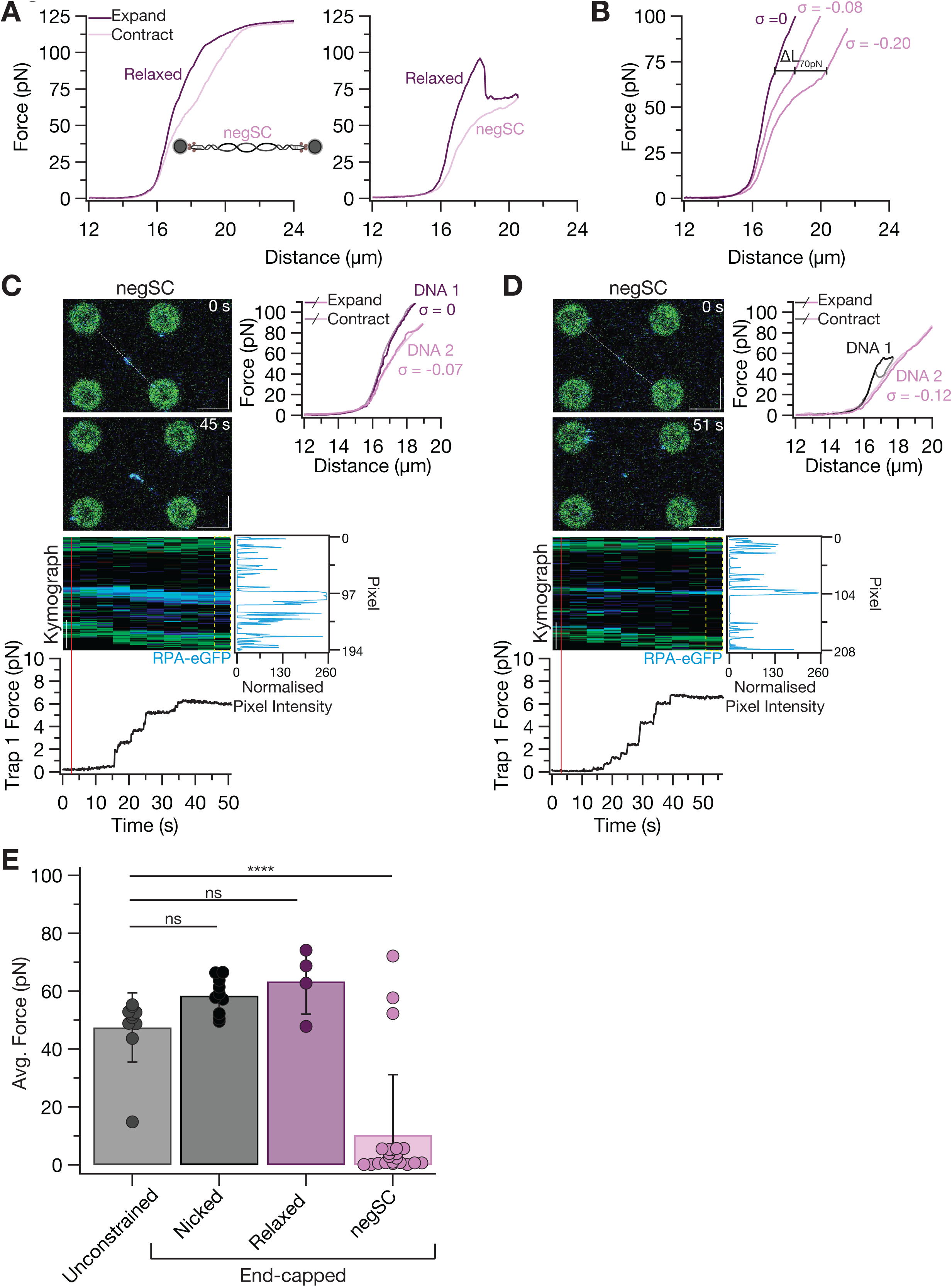
Generating braids with negatively supercoiled DNA results in ssDNA bubbles at low force. **(A)** (left) Example of an FE curve generated by a DNA that begins relaxed (dark purple) and becomes negatively supercoiled (negSC) (light purple) after extending to high force and contracting. (right) Example of an FE curve generated by a relaxed DNA that becomes negatively supercoiled during the extension. **(B)** Examples of two stable negSC DNAs with different supercoiling densities (σ = -0.08 and -0.20) calculated from their extension at 70 pN. ^28,47^ **(C-D)** Similar to Figure 1E-F. FE curves recorded in channel 3 show the type of DNA between each bead pair and the calculated supercoiling density (σ) if the DNA is a constrained molecule. Black denotes a nicked DNA, dark purple for relaxed, and light purple for negatively supercoiled. Experiments were done in the presence of 5 nM RPA-eGFP. **(E)** Average force in trap 1 and trap 2 required for RPA binding at braid for different types of DNA. Braids with negatively supercoiled DNA were generated with at least one DNA with an average σ of 0.08 ± 0.07. Mean force reported with SD (error bars): Unconstrained = 47.47 pN ± 11.94 pN (n = 10), Nicked = 58.48 pN ± 6.23 pN (n = 10), Relaxed = 63.38 pN ± 11.34 pN (n = 4), negSC = 10.38 pN ± 20.82 pN (n = 22) **** denotes a p-value < 0.0001 calculated by a one-way ANOVA and Tukey HSD/Tukey Kramer test. Scale bars = 5 μm.

Negative supercoiling (σ = -0.07 and -0.12) in just one of the DNAs dramatically reduced the force required for RPA recruitment to the DNA-DNA crossing to forces near 0 pN (Fig. 2C-D) especially when compared to relaxed constrained molecules (Fig. 1F). Notably, this effect was caused by any level of supercoiling above a critical threshold of σ ∼ -0.03 (Fig. S1B). The majority of the braids (19 out of n = 22) composed of DNA with at least one of the DNAs having σ of -0.02 – -0.32 had RPA binding at the DNA-DNA braid at the drastically reduced force (Fig. S1B-C). Taken together, comparison across the different substrates tested shows that negative supercoiling significantly lowers the force required to expose single-stranded DNA and recruit RPA at the crossing (Fig. 2E).

### Catenane geometry lowers force threshold for ssDNA formation in negatively supercoiled DNA

We next asked whether the strong ssDNA exposure observed in negatively supercoiled braids reflected a general destabilisation of negatively supercoiled DNA or whether the DNA-DNA crossing itself preferentially promoted local melting. To distinguish between these possibilities, we used dual-trap optical tweezers to compare RPA recruitment to individual DNA molecules subjected to increasing tension that were either relaxed (Fig. 3A) or negatively supercoiled to the same extent as those we used above to form DNA braids (Fig. 3B-C, S2A).

**Figure 3.**
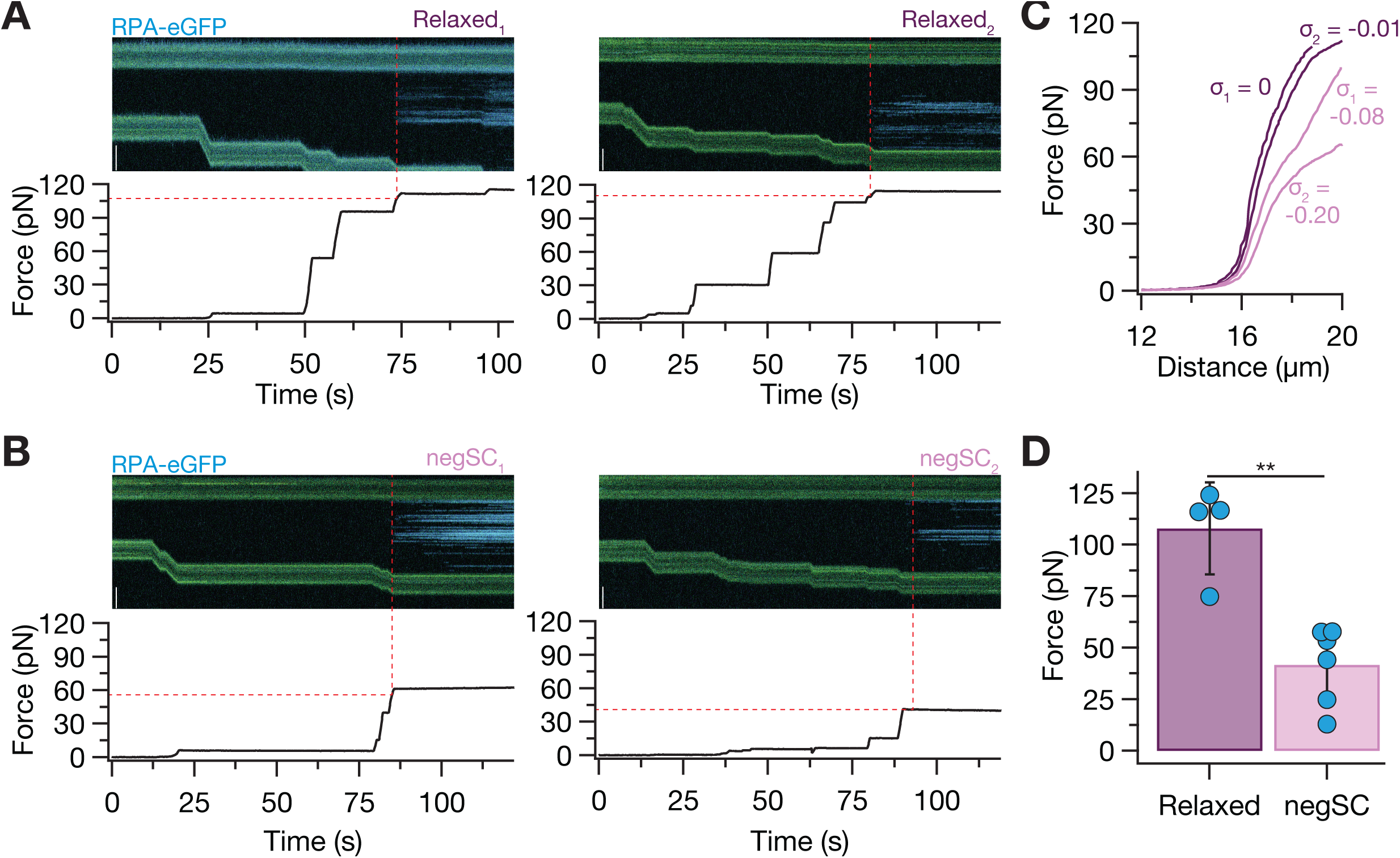
RPA binds to braided supercoiled DNA at lower forces than on linear supercoiled DNA. **(A)** Two representative kymographs of a single relaxed DNA extended in a channel containing 5 nM RPA-eGFP with the corresponding force. Dotted red lines indicate the force at which RPA bound. Scale bars = 5 μm. **(B)** Same as panel A with two examples of negatively supercoiled DNAs of different supercoiling densities (σ = -0.08 and -0.20). Scale bars = 5 μm. **(C)** FE curves of the DNAs shown in panels A and B and their calculated supercoiling densities. **(D)** Comparison the force at which RPA bound to a single relaxed or supercoiled DNA. Mean force in trap 2 reported with SD (error bars): Relaxed = 107.88 pN ± 22.33 pN (n = 4), negSC = 41.74 pN ± 18.84 pN (n = 6). ** denotes a two-tailed p-value of 0.0010 as calculated by an un-paired *t* test.

RPA recruitment depended strongly on both negative supercoiling and applied tension (Fig. 3D). Negatively supercoiled molecules (σ = -0.02 – -0.35) were susceptible to strand separation, with RPA recruitment occurring at tensions of ∼13-58 pN in 125 mM potassium glutamate, whereas relaxed molecules experienced strand separation at 75-124 pN (Fig. 3A-B, D). While the amount of RPA binding to ssDNA will depend on the amount of supercoiling and salt in the experimental buffer, these results are consistent with previous observations that negative supercoiling and mechanical tension cooperate to destabilise duplex DNA ^27,28^. However, the presence of negatively supercoiled DNA in a braid clearly reduced the force threshold for RPA recruitment. The force needed to allow RPA binding on a single negatively supercoiled DNA molecule was correlated with the amount of supercoiling (Fig S2B); whereas any amount of supercoiling was enough to reduce the force needed for RPA binding specifically at the crossing of braided DNA (Fig. 2E, S1B).

Together, these observations identify DNA-DNA crossings as favoured sites for duplex destabilisation within negatively supercoiled DNA. We conclude that negative supercoiling sensitises the duplex to strand separation, while the geometry caused by braiding the DNA substantially lowers the mechanical threshold at which ssDNA becomes exposed. This suggests that torsional stress might convert an otherwise duplex DNA entanglement into a substrate accessible to the single-strand passage mechanism of TOP3α.

### Negative supercoiling enables TRR to decatenate DNA braids

Our observation that negative supercoiling promotes local ssDNA exposure specifically at DNA crossings suggests a mechanism by which an apparently duplex DNA entanglement might become accessible to human TOP3α. First, we purified the human TOP3α–RMI1–RMI2 complex (TRR) and labelled TOP3α with Alexa Fluor 555 (TRR-AF555) (Fig. S2C). The activity of labelled TRR was confirmed using DNA relaxation assays (Fig. S2D). Next, we investigated TRR recruitment to negatively supercoiled molecules (σ = -0.04 – -0.08). TRR recruitment to single molecules that were either relaxed or negatively supercoiled closely paralleled that of RPA (Fig. S2B, E-H), similar to that reported recently ^29^. Thus, recruitment of TRR (TOP3α) correlates with conditions that expose ssDNA.

To test whether TRR could decatenate DNA braids we first examined DNA braids formed with end-capped DNA molecules with nicks, rendering them unconstrained. Under these conditions, detectable RPA recruitment to the crossing required high forces > 60 pN (Fig. 2E). Braids were held under increasing force in the presence of 5 nM RPA until binding occurred at the junction (Fig. 4A, S3A) and were subsequently transferred into a channel containing 2 nM TRR while maintaining the applied force. No detectable decatenation was observed under these conditions (Fig. 4A, S3A) and often one of the DNA molecules forming the braid broke within the sampling time of 2 minutes after entering the channel containing TRR (5 out of n = 7) (Fig. 4A, S3A-B). This breakage is consistent with previous observations reporting that at forces above ∼50 pN the non-covalent interaction between the TRR complex and the cleaved 3′ end of the ssDNA is disrupted ^30^, thus converting the transient TOP3α cleavage intermediate into a permanent break. Thus, although high mechanical tension can expose ssDNA within the braid containing relaxed DNAs, at this tension the substrates were not productively resolved by TRR (Fig. 4A-B).

**Figure 4.**
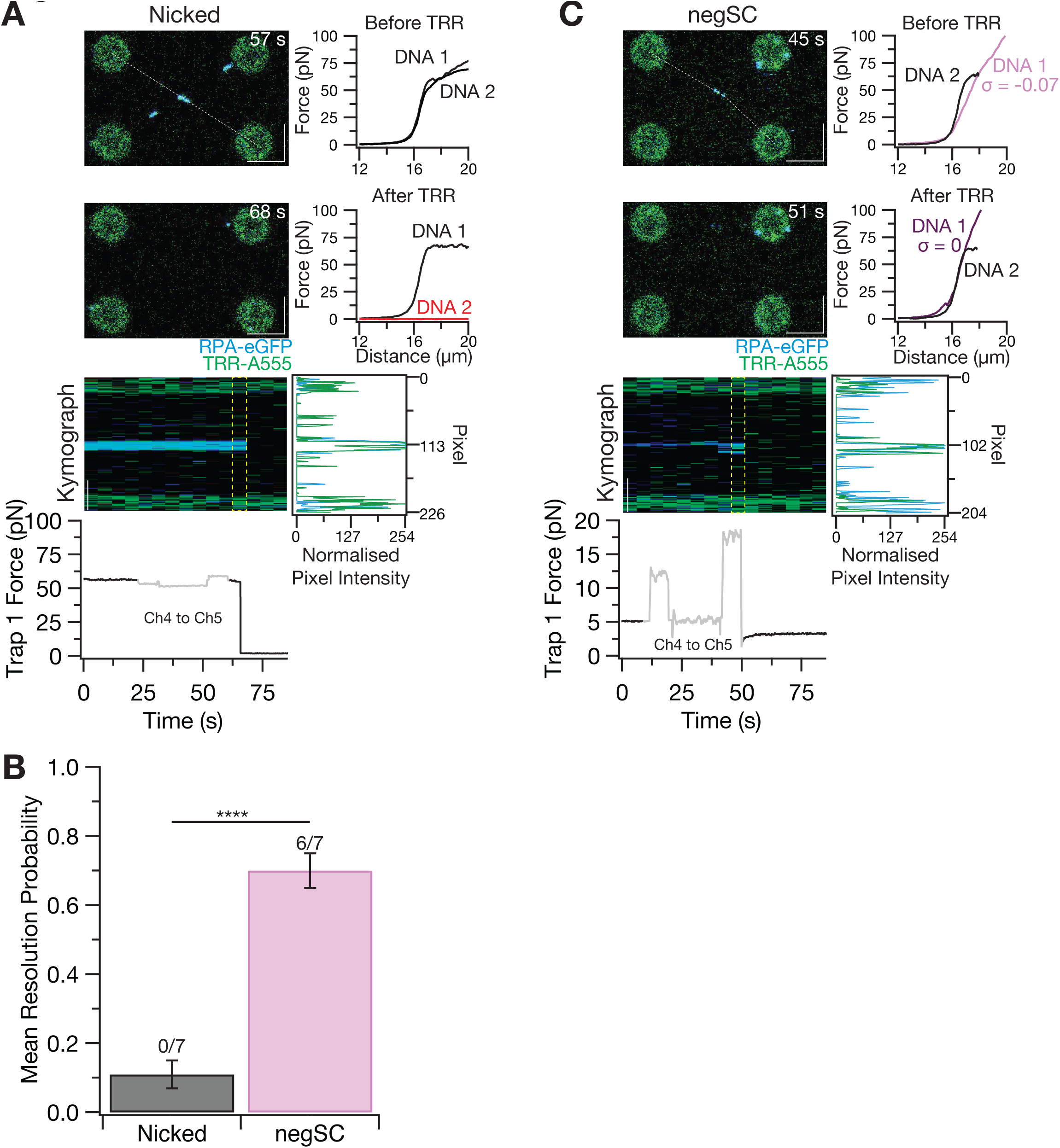
TRR decatenates braids made with negatively supercoiled DNA. **(A)** Examples of frames from a 2-D confocal scan of a braid made with two end-capped, nicked DNAs that were first stretched in channel 4 by moving beads 2 and 3 in the x-axis with only 5 nM RPA-eGFP present until RPA bound to the braid (bead 1 force = ∼ 60 pN). A new scan was started in the RPA channel after RPA was detected at the braid and continued while the braid was moved to a channel that contained 2 nM TRR. The selected frames show the braid before and after fluorescence signal at the braid was lost, with a kymograph of the whole scan (acquired along the white dashed line) and the corresponding force showing the move from channel 4 to 5 in the grey box. The FE curves were taken before (in channel 3) and after (in channel 5) the experiment and confirm that the drop in force was caused by DNA 2 breaking. The normalised blue and green pixel intensity of the frame in the kymograph denoted with the dotted yellow line shows a peak at the centre of the braid. Scale bars = 5 μm. **(B)** Mean resolution probability in a 2-minute sampling time of TRR decatenation of braids made with either two end capped, nicked DNAs at or at least one negatively supercoiled DNA after RPA binding. Error bars indicate standard error. **** denotes a two-tailed p-value < 0.0001 as calculated by an un-paired *t* test. **(C)** Similar to panel A but the braid is made of one negatively supercoiled DNA (DNA 1), σ = -0.07, and one nicked DNA (DNA 2). Braid was set to 5pN (on bead 1 and 4) after RPA binding occurred in channel 4 and before moving to channel 5. FE curves show two intact DNAs before (in channel 3) and after the loss of fluorescence signal at the braid (in channel 5), indicating decatenation of the braid (as well as relaxation of the supercoiled DNA). Scale bars = 5 μm.

We next examined braids in which one or both DNA molecules were negatively supercoiled. Similar to the braids made with nicked DNA, braids containing negative supercoiling were incubated with 5 nM RPA at 0 pN and stretched until binding was observed at the junction. For optimal imaging, the braids were then held at 5 pN before being moved to the channel containing TRR, regardless of the force that RPA bound. These braids were resolved with high efficiency (Fig. 4B-C, S3C), with decatenation occurring within 10s of seconds following exposure to 2 nM TRR (6 out of n = 7) (S3B). Additionally, the supercoiled DNA was relaxed after the decatenation experiment, consistent with the mechanism described recently ^29^ (Fig. 4C, S3C).

Importantly, decatenation of negatively supercoiled braids occurred at 5 pN (Fig. 4C, S3C), corresponding closely to the force regime in which RPA first detected ssDNA at the crossing of highly negatively supercoiled braids (Fig. 2E). Thus, the ability of TRR to resolve DNA braids correlates with the appearance of locally melted DNA at the junction. These results suggest that negative supercoiling converts an otherwise refractory duplex DNA crossing into an efficient substrate for TRR.

### RPA and RMI2 stimulate TOP3α-mediated decatenation

Previous biochemical studies with the yeast Sgs1–Top3–Rmi1 complex showed that RPA strongly stimulates Top3-mediated catenation and decatenation reactions ^13^. Because RPA was present and highly correlated with braid resolution by TRR when one or both DNAs were negatively supercoiled (Fig. 4B, S3B), we asked whether RPA was necessary for this activity.

To this end, we generated negatively supercoiled braids that were maintained at ∼5 pN, corresponding to conditions in which RPA readily binds the junction (Fig. 2E). Rather than pre-incubating these substrates with RPA, braids were transferred directly into a channel containing 2 nM TRR. TRR remained capable of resolving these negatively supercoiled braids in the complete absence of RPA (2 out of n = 6) (Fig. 5A, S3D-E), demonstrating that RPA is not essential for the reaction. However, the probability of decatenation was nevertheless substantially reduced compared with reactions in which RPA had first been allowed to bind the crossing (Fig. 5B). Furthermore, TRR binding was not observed at the braid at 5 pN in any of the experiments (0 out of n = 6) (Fig. 5A). Thus, RPA strongly stimulates, but is not required for, TRR-mediated decatenation. These observations are consistent with a model in which the critical function of RPA is to trap transiently melted ssDNA bubbles at negatively supercoiled crossings in the open state. By binding the exposed single strands and preventing their reannealing, RPA greatly extends the lifetime of the bubble, and hence the window during which TRR can engage it for strand passage.

**Figure 5.**
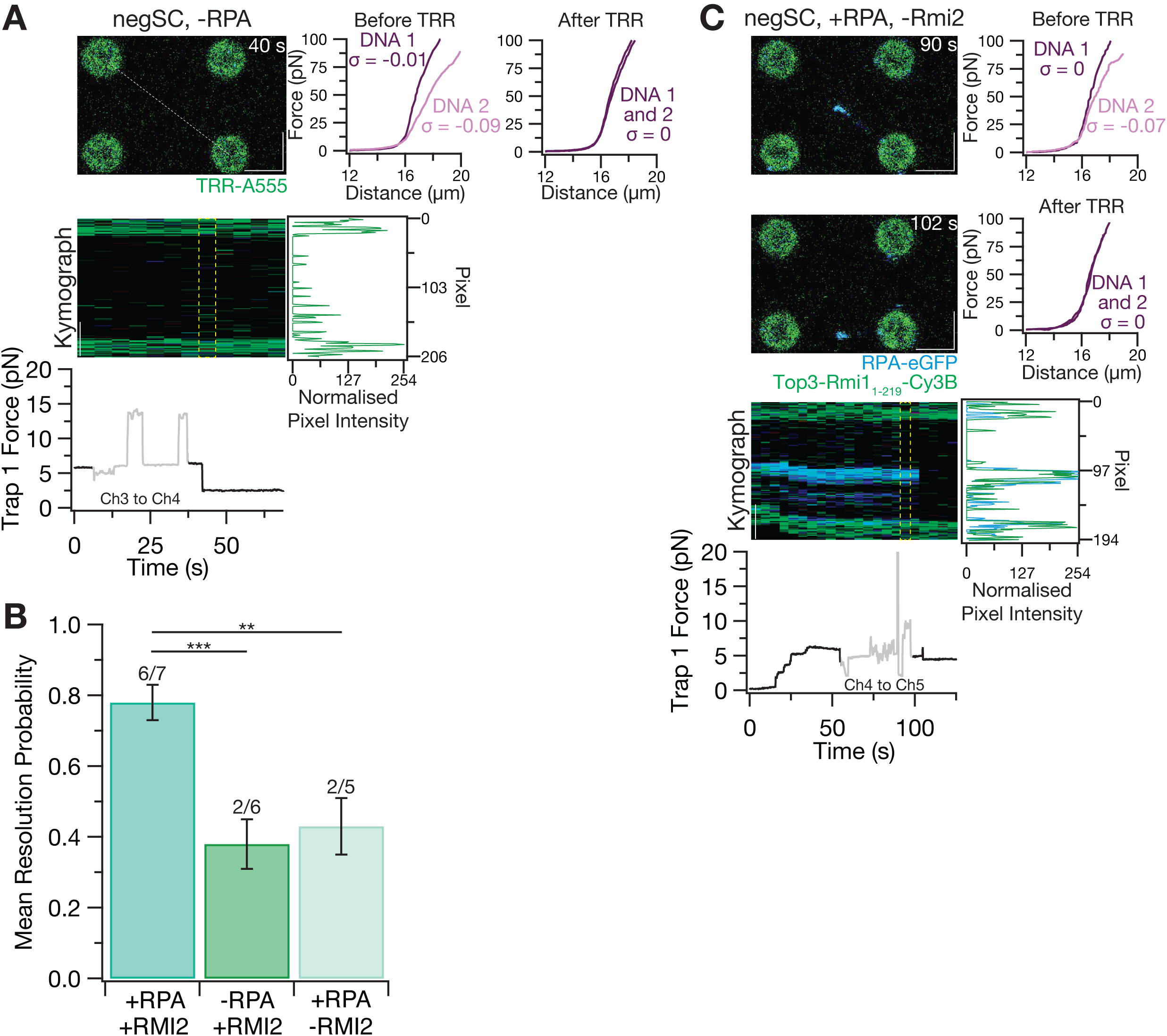
Dependency of RPA and RMI2 on TRR decatenation of negatively supercoiled braid. **(A)** Example of decatenation experiment performed in the absence of RPA. Examples of frames from a 2-D confocal scan of a braid made with one relaxed DNA (DNA 1) and one negatively supercoiled DNA (DNA 2), σ = -0.09, that was set to 5 pN (on bead 1 and 4) in channel 3 before being moved to channel 4 containing 2 nM TRR-A555. Kymograph and corresponding force show the move from channel 3 to channel 4 in the grey box. FE curves before (in channel 3) and after (in channel 4) the experiment show two intact DNA, indicating decatenation of the braid (and relaxation of supercoiling). The green channel pixel intensity plot shows the normalised intensity in the green channel along the braid at the time of decatenation and the lack of any TRR-A555 binding observed at the time of decatenation. Scale bars = 5 μm. **(B)** Mean resolution probability in a 2-minute sampling time of TRR on negatively supercoiled braids at 5 pN ± RPA and ± RMI2 shown with standard error. Two independent unpaired *t* tests were conducted comparing TRR ± RPA (two-tailed p-value = 0.0004) and TRR ± RMI2 (two-tailed p-value = 0.0023). **(C)** Examples of frames from a 2-D confocal scan of a braid made with one relaxed DNA (DNA 1) and one negatively supercoiled DNA (DNA 2), σ = -0.07. Braid was set to 5pN (on bead 1 and 4) after RPA binding in channel 4 with 5 nM RPA-eGFP and before moving to channel 5 containing 2 nM TOP3α-RMI1_1-219_. The scan was started in channel 4 and continued while the braid was moved to channel 5. The selected frames show the braid before and after fluorescence signal at the braid was lost, with a kymograph of the whole scan and corresponding force showing the move from channel 4 to channel 5 in the grey box. FE curves before (in channel 3) and after (in channel 5) the experiment and show two intact DNA, indicating decatenation of the braid (and relaxation of supercoiling). The blue and green pixel intensity of the frame in the kymograph denoted with the dotted yellow line shows a peak at the centre of the braid. Scale bars = 5 μm.

We next investigated the contribution of RMI2 to TRR-mediated decatenation. RMI2 associates with the C-terminal OB-fold of RMI1 to form the RMI1C–RMI2 module and has been proposed to provide a regulatory and stabilising layer within the TOP3α–RMI1–RMI2 complex ^31,32^. In contrast, the N-terminal region of RMI1 interacts directly with TOP3α and is sufficient, together with TOP3α and BLM, to support double Holliday junction dissolution in vitro ^31,33^.

To determine whether the RMI1C–RMI2 module was required for decatenation of negatively supercoiled DNA braids, we compared the complete TRR complex with a minimal TOP3α–RMI1 complex containing RMI1 residues 1–219 (TOP3α-RMI1_1-219_). We purified and fluorescently labelled the minimal TOP3α–RMI1_1-21_ complex which showed activity in relaxation assays (Fig. S2D). This complex retained clear decatenation activity against negatively supercoiled DNA braids (Fig. 5B-C), demonstrating that neither RMI2 nor the C-terminal region of RMI1 is essential for the reaction. Decatenation efficiency, however, was reduced by approximately 50% relative to the complete TRR complex (Fig. 5B, S3D-E). Thus, TOP3α together with the N-terminal region of RMI1 is capable of resolving negatively supercoiled DNA braids, whereas the RMI1C–RMI2 module enhances the efficiency of the reaction.

### TRR resolves DNA braids by passing duplex DNA through a single-stranded region

The requirement for ssDNA at braid crossings implies that TOP3α resolves these structures by the canonical type IA mechanism, in which one strand is transiently cleaved, and a second DNA segment is transported through the resulting opening. To test this directly, we used a substrate in which the single-stranded region was defined in advance rather than generated by supercoiling-induced duplex melting, so that the passed segment and the gated single strand could be unambiguously assigned.

We therefore intertwined one λ-DNA molecule containing a defined ssDNA gap with a second, fully double-stranded DNA (dsDNA) molecule. A hybrid dsDNA-ssDNA-dsDNA substrate was generated by stretching end-capped DNA molecules that happen to have two nicks on the same DNA strand to ∼100 pN (Fig. 6A) followed by force-induced unpeeling of the intervening sequence ²⁸ to form a ‘gapped’ molecule, based on the protocol from Belan *et al.* 2021 ^34^ (Fig. 6B). Gap formation was confirmed from the characteristic FE behaviour of the resulting molecules (Fig. 6C). Braids formed with one gapped DNA and one dsDNA were then introduced to a channel containing both 5 nM RPA-eGFP and 2 nM TRR-A555 (Fig. 6D)

**Figure 6.**
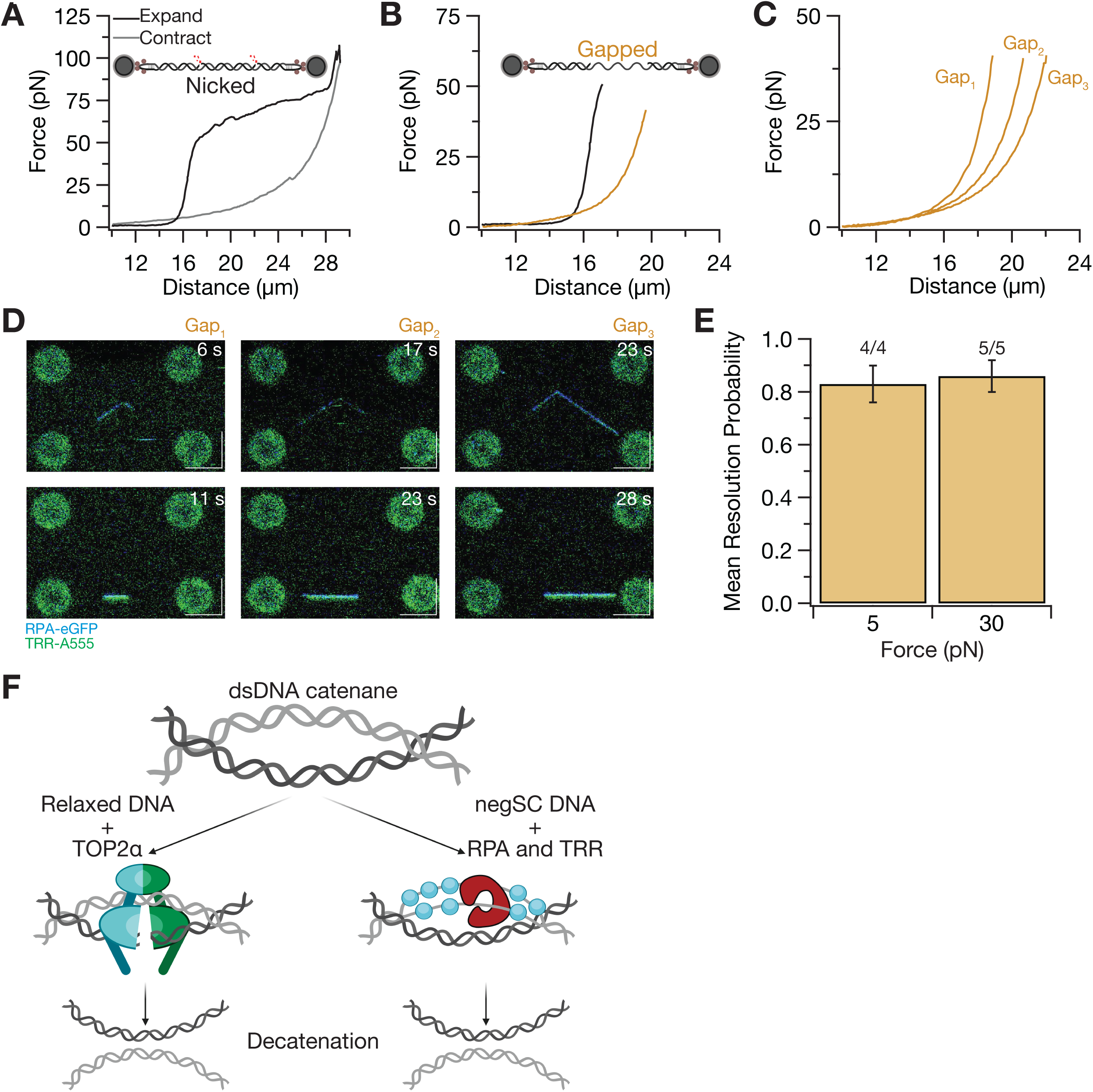
TRR is able to pass a duplex DNA through a single stranded gap. **(A)** FE curve of an end-capped DNA with more than one nick on the same strand stretched to ∼100 pN and contracted to form DNA with a single stranded gap. **(B)** FE curves of the DNA in panel A before (black) and after (yellow) formation of the single stranded gap. The presence of the gap is confirmed by the extension at a given force compared to that of the original DNA. **(C)** Examples of different length gaps formed from nicked DNA. **(D)** Frames from 2-D confocal scans of braids formed with the gapped DNA from panel C and a non-gapped DNA entering a channel containing both 5 nM RPA-eGFP and 2 nM TRR and becoming resolved. Examples shown were conducted at 30 pN. Scale bars = 5 μm. **(E)** Mean resolution probability in a 2-minute sampling time of one gapped DNA and one dsDNA by TRR at 5 and 30 pN. **(F)** Proposed model of dsDNA catenane resolution pathways. In the case of relaxed DNA catenanes, TOP2α is the primary resolving enzyme. In the case of negatively supercoiled DNA catenanes, RPA is able to bind and stabilise the single stranded bubbles localising at the crossover, allowing TRR to resolve the intertwine.

In this configuration the only single-stranded region in the braid lies within one of the two molecules, while its partner remains fully duplex throughout (Fig. 6D). TRR catalysed efficient resolution of these braids when held at both 5pN and 30pN (Fig. 6E). Because no ssDNA is generated (and observed) in the unconstrained duplex partner under these conditions, resolution can only have occurred by passage of an intact duplex segment through the single-stranded gap of the gapped molecule (Fig. 6D). This establishes that TOP3α decatenates braided DNA by type IA strand passage, and that a pre-existing single-stranded region within one partner is sufficient to license the reaction.

Collectively, our results reveal how the physical state of DNA can determine the pathway available for resolving DNA entanglements. Negative supercoiling promotes local duplex melting specifically at DNA-DNA crossings, generating ssDNA that can be captured by RPA and engaged by TRR (Fig. 6F). This converts an otherwise inaccessible duplex DNA crossing into a substrate for type IA strand passage, in which a duplex segment is transported through the single-stranded region of its partner, providing a mechanism by which torsional stress can direct the resolution of DNA linkages towards a single-strand-dependent topoisomerase pathway.

## Discussion

The resolution of DNA entanglements is generally associated with type II topoisomerases, whose ability to transport an intact DNA duplex through a transient double-stranded break provides a direct mechanism for removing inter-DNA linkages ^15^. Type IA topoisomerases face a fundamentally different constraint, where strand passage occurs through a transient break in ssDNA ^15^ and therefore requires access to ssDNA within the substrate. To this end, bacterial and yeast Top3 enzymes require the helicase activity of Sgs1 to promote decatenation reactions involving apparently duplex DNA molecules ^11,13^. Notably, Sgs1 can unwind duplex DNA at internal sites, without requiring a free end ^13^ thus generating the single-stranded regions that Top3 requires. Here, we identify a physical mechanism capable of bypassing the requirement of a helicase. We show that negative supercoiling promotes local duplex melting specifically at DNA-DNA crossings and thereby enables human TRR to resolve a double-stranded braid. Thus, the topological state of DNA can determine not only the geometry of an entanglement, but also its biochemical accessibility to different classes of topoisomerase.

A central observation of our study is that DNA crossings are particularly susceptible to supercoiling-dependent duplex destabilisation (Fig. 2C-E). Negative supercoiling stores torsional energy within DNA and local disruption of base pairing is one mechanism through which this stress can be accommodated ^27,28^. Consistent with this, introducing negative supercoiling reduced the force required to expose ssDNA in individual DNA molecules (Fig. 3D, S2B). However, this effect was substantially amplified when negatively supercoiled DNA was incorporated into a braid (Fig. 2E, S1B). Whereas detectable ssDNA formation on an isolated highly negatively supercoiled DNA molecule required ∼40 pN of applied tension under the buffer conditions used here (Fig. 3D), RPA accumulated at the crossing of negatively supercoiled braids at forces under ∼1 pN (Fig. 2E). The geometry of a DNA-DNA crossing therefore promotes duplex destabilisation when combined with negative torsional stress.

The physical basis for this enhanced susceptibility remains to be established. DNA-DNA crossings impose local geometrical constraints and could influence DNA bending, steric interactions and the partitioning of torsional stress between twist, writhe and alternative DNA conformations. Any combination of these effects could reduce the energetic cost of disrupting base pairing at the crossing. That base pairing is indeed locally disrupted at such sites is supported by earlier intercalator-based force measurements on braided DNA, where unconstrained molecules appeared to fuse at forces of approximately 40 pN, an effect attributed to a structural interaction between locally denatured regions at the point of entanglement ^21^. Importantly, our experiments do not require the junction to contain a stable or extended region of ssDNA. Instead, we favour a model in which negative supercoiling increases the probability and/or lifetime of transiently melted states at the crossing. In our experiments RPA captures these states, providing a sensitive experimental readout of their formation (Fig. 2C-D). The striking reduction in the force threshold for RPA recruitment when negatively supercoiled DNA is incorporated into a braid suggests that crossing geometry itself strongly favours this transition. Determining the dimensions, lifetime and precise structure of the melted intermediate will require approaches capable of probing DNA structure at greater spatial and temporal resolution.

Our data show that this local structural transition has a profound consequence for topoisomerase specificity. Relaxed duplex braids were refractory to TRR-mediated resolution (Fig. 4A, C, S3A), whereas negatively supercoiled braids were rapidly and efficiently decatenated (Fig. 4B-C, S3C). In the former condition, TRR was more likely to break one of the DNAs than resolve the braid into two intact DNAs (Fig. S3B). Moreover, the conditions that promoted TRR-mediated decatenation closely corresponded to those that generated detectable ssDNA at the junction. These observations strongly support a mechanism in which negative supercoiling changes the physical nature of the DNA crossing, generating the single-stranded substrate required for type IA strand passage. Negative supercoiling therefore does not simply enhance TOP3α recruitment or globally increase DNA accessibility. Rather, torsional stress converts a dsDNA-dsDNA crossing into a junction containing sufficient ssDNA character to support TOP3α catalysis.

This mechanism provides a possible explanation for previous observations that type IA topoisomerases can catalyse reactions involving apparently duplex DNA substrates. In *E. coli*, Topo III can promote decatenation of replicating daughter DNA molecules, an activity that is strongly stimulated by the RecQ helicase ^11,12^. Given that *E. coli* DNA is maintained in a negatively supercoiled state *in vivo* ^35,36^ through the activity of DNA gyrase ^37^, our findings raise the possibility that these apparently duplex substrates in fact contain localised regions of underwound or single-stranded character arising from this physiological negative supercoiling, analogous to the junction ssDNA we observe in negatively supercoiled braids in vitro. Similarly, the budding yeast Sgs1–Top3–Rmi1 complex promotes catenation and decatenation of duplex plasmid substrates *in vitro*, reactions that require both Sgs1 and RPA^13^. In contrast, studies of the human complex have focused on TOP3α-dependent relaxation of negatively supercoiled DNA^8^, while catenation or decatenation of duplex DNA has not, to our knowledge, been assayed for this complex. The decatenation activity reported for human TOP3α is instead confined to single-stranded catenanes ^30^. Our observations suggest that apparently duplex topological substrates do not require helicase activity to unwind dsDNA from internal sites. Instead, torsional stress and crossing geometry can cooperate to create transient ssDNA precisely at the topological junction, thereby providing an entry point for type IA topoisomerases.

The stimulatory effect of RPA is consistent with this model. RPA was not essential for TOP3α-mediated decatenation (Fig. 5A), demonstrating that negative supercoiling itself is sufficient to generate a productive substrate. Nevertheless, the efficiency of braid resolution was substantially enhanced by RPA (Fig. 5B). A straightforward interpretation is that RPA captures transiently exposed ssDNA at the crossing and limits its reannealing, increasing the lifetime of the substrate available for TOP3α. Such a mechanism could explain the strong stimulation of Top3-dependent catenation and decatenation by RPA observed with the yeast Sgs1–Top3–Rmi1 complex ^13^. More generally, it illustrates how ssDNA-binding proteins can facilitate, rather than simply compete with type IA topoisomerases, by stabilising transient DNA structures required for their activity.

The contribution of RMI1–RMI2 further indicates that substrate formation and catalytic efficiency represent distinct levels of regulation. The TOP3α–RMI1_1-219_ complex retained the ability to resolve negatively supercoiled DNA braids (Fig. 5C), demonstrating that RMI2 and the C-terminal region of RMI1 are not strictly required for decatenation. Their absence nevertheless substantially reduced reaction efficiency (Fig. 5B). This is consistent with structural and biochemical studies indicating that RMI1 and RMI2 regulate the TOP3α strand-passage and stabilise catalytic intermediates ^31–33^. Negative supercoiling therefore creates the ssDNA-containing crossing required for TOP3α engagement, whereas RPA and RMI1–RMI2 increase the probability that this intermediate undergoes productive strand passage.

Our findings suggest that DNA topology could contribute to determining which topoisomerase pathway acts on a particular DNA entanglement in cells (Fig. 6F). Replication and transcription continuously generate torsional stress, while chromosomal DNA is organised into topologically constrained domains in which free rotation is restricted. Replication in particular creates a dynamic topological environment in which positive supercoiling ahead of replication forks and intertwining of newly replicated DNA behind them are mechanically coupled. Local redistribution of torsional stress could therefore generate regions of negative supercoiling within entangled DNA and promote transient strand separation at particular crossings. Our experiments do not establish the magnitude or distribution of such torsional stress *in vivo*, but they reveal that, once present, negative supercoiling can dramatically alter the accessibility of a DNA linkage to TOP3α.

Torsional stress is not the only route to a single-stranded region within entangled DNA. The gapped substrate used here has a direct physiological counterpart: single-stranded gaps are generated behind replication forks, through discontinuous lagging-strand synthesis and through repriming of the leading strand, and persist until they are filled by post-replicative repair ^38^. A molecule carrying such a gap that becomes intertwined with its sister would present TOP3α with the configuration we have reconstituted (Fig. 6D), namely a duplex segment that can be passed through a pre-existing single-stranded region with no requirement for local melting. TOP3α might therefore contribute not only to the resolution of entanglements between two intact duplexes, where ssDNA must first be generated at the crossing, but also to the removal of linkages involving DNA that is already gapped, during post-replicative gap repair, template switching and the processing of recombination intermediates, in which ssDNA is an intrinsic feature of the structures involved. Because ssDNA is required only for strand cleavage, and the segment transported through the resulting break may be single-stranded or duplex ^32^, one gapped partner suffices for unlinking. Such an activity would extend the established ssDNA-dependent functions of TOP3α, best characterised in the dissolution of double Holliday junctions, to topological linkages that arise while replication-associated gaps remain unfilled, and would provide a mechanistic basis for the proposal that bacterial topoisomerase III acts at the replication fork, using gaps in the nascent strands as entry points, to remove precatenanes between partially replicated sister chromosomes ^39^.

RecQ-family helicases provide an additional route through which the ssDNA required for TOP3α activity could be generated. The functional cooperation between RecQ helicases and Top3 enzymes is evolutionarily conserved. *E. coli* RecQ strongly stimulates Top3-mediated decatenation ^12^, Sgs1 functions with Top3–Rmi1 in budding yeast, and human BLM forms the BTRR complex with TOP3α-RMI1-RMI2. BLM-mediated unwinding could generate or enlarge ssDNA at an entangled DNA junction and thereby provide TOP3α with the substrate required for strand passage. Importantly, BLM-dependent unwinding and supercoiling-induced melting need not represent alternative mechanisms. Negative torsional stress could lower the energetic barrier to initial duplex opening at a DNA crossing, after which BLM could enlarge or stabilise the melted region. RPA could then capture the exposed strands and limit their reannealing, extending the lifetime of the substrate available to TOP3α. In this view, TOP3α does not necessarily recognise a particular class of DNA entanglement *per se*; instead, multiple processes can generate the ssDNA-containing crossing that TOP3α recognises and resolves.

Such a mechanism may be particularly relevant to ultrafine anaphase bridges (UFBs). UFBs are thin DNA threads connecting segregating sister chromatids during anaphase and can arise from unresolved DNA entanglements, incompletely replicated regions and recombination intermediates ^40,41^. In contrast to conventional chromatin bridges, UFBs are largely devoid of canonical nucleosomal chromatin and are decorated by PICH, BLM, and other factors involved in their processing. TOP3α and its partners RMI1 and RMI2 are components of this UFB-associated machinery, placing the BTRR complex at highly stretched and potentially topologically constrained DNA linkages during chromosome segregation ^40,41^.

PICH could contribute to establishing the physical DNA environment required for resolution of these structures ^42^. PICH preferentially associates with DNA under tension and has been implicated in maintaining UFBs in a nucleosome-depleted state ^43^. Single-molecule studies further indicate that PICH can promote force-dependent nucleosome unwrapping and remodelling ^44^. Such chromatin remodelling may have topological consequences in addition to exposing DNA to repair and resolution factors. DNA wrapping around nucleosomes constrains linking number; consequently, displacement of nucleosomes from DNA whose rotational freedom remains restricted could redistribute the linking difference previously accommodated by nucleosomal wrapping into the exposed DNA. Depending on the topological boundary conditions of the bridge, this could generate or redistribute torsional stress and potentially favour negative supercoiling within regions of UFB DNA.

Our results therefore suggest a speculative but testable model for how the unusual physical organisation of UFBs could facilitate TOP3α-dependent resolution. As sister chromatids segregate, increasing tension across an unresolved DNA linkage would favour PICH association and could promote PICH-dependent nucleosome remodelling, exposing the underlying DNA and altering the distribution of torsional stress within the bridge. If this generates or concentrates negative torsional stress, DNA-DNA crossings within the entanglement would become increasingly susceptible to local duplex melting. BLM could further unwind these destabilised regions, while RPA could capture and stabilise the resulting ssDNA. Together, these processes would generate precisely the type of ssDNA-containing DNA crossing that our experiments identify as an efficient substrate for TRR-mediated strand passage.

This model also provides a framework for considering how TOP2α and TOP3α might contribute differently to the resolution of persistent anaphase DNA linkages (Fig. 6F). TOP2α is intrinsically capable of resolving intact duplex–duplex catenations and is therefore ideally suited to conventional DNA decatenation. TOP3α, by contrast, would become effective when the local physical environment of an entanglement favours duplex opening. Progressive stretching, chromatin remodelling, torsional stress and BLM activity could therefore change the biochemical identity of a persistent DNA crossing during chromosome segregation. A crossing that initially constitutes a substrate primarily for TOP2α could acquire ssDNA character and thereby become accessible to TOP3α. TOP3α may also act on such linkages indirectly: PICH and TOP3α have been proposed to function together as a reverse gyrase, in which PICH extrudes supercoiled loops whose negative supercoils TOP3α selectively relaxes, leaving positive supercoiling in the surrounding DNA ^45^ and generating an optimal substrate for TOP2α-mediated disjunction of sister centromeres ^46^. The two routes are not mutually exclusive, and both point to the same underlying principle: the topological state of a persistent crossing determines which enzyme can act on it, and that state is itself actively remodelled. Such pathway flexibility may be important for persistent or structurally complex DNA linkages that remain unresolved as cells enter anaphase, particularly because UFBs are thought to endure extreme forces and our experiments show TRR decatenation is still viable at ∼30 pN (Fig. 6E).

Several aspects of this model remain to be tested. Our experiments establish that negative supercoiling promotes local ssDNA formation at DNA crossings and that this enables TOP3α-mediated decatenation *in vitro*, but they do not establish that negatively supercoiled crossings constitute physiological TOP3α substrates in cells. Nor do they define the amount or sign of torsional stress present within UFBs. In particular, although nucleosome displacement from topologically constrained DNA has the potential to redistribute linking number, whether PICH-dependent chromatin remodelling generates negative torsional stress45 within UFBs remains unknown. Direct measurements of DNA topology within UFBs, and experiments determining whether manipulation of torsional stress influences BTRR-dependent bridge resolution, will be required to test this model.

More broadly, our findings reveal how DNA mechanics can determine enzymatic substrate specificity. An entanglement that is inaccessible to TOP3α when formed between relaxed DNA duplexes becomes an efficient substrate when negative supercoiling promotes local strand separation at the crossing. Torsional energy stored within DNA can therefore be converted into a local structural transition that enables an alternative strand-passage mechanism. We propose that DNA supercoiling can act as a physical determinant of topoisomerase pathway choice, coupling the mechanical and topological state of DNA to the enzymatic resolution of DNA entanglements.

## Materials and Methods

### Protein expression constructs

| Construct | Subunits | Plasmid | Expression system | Tags | Source |
| --- | --- | --- | --- | --- | --- |
| *Hs*TRR | TOP3α<sub>FL</sub>,<br>RMI1<sub>FL</sub>,<br>RMI2<sub>FL</sub> | pBIG1a<br>(Addgene<br>80611) | *Sf9*-<br>Baculovirus | TOP3α-<br>HRV 3cs-<br>ybbr-TEV-Strep-Tag II | This study |
| *Hs*TOP3-RMI1<sub>1-219</sub> | TOP3α<sub>FL</sub>,<br>RMI1<sub>1-219</sub> | pBIG1a<br>(Addgene<br>80611) | *Sf9*-<br>Baculovirus | TOP3α-<br>HRV 3cs-<br>ybbr-TEV-Strep-Tag II | This study |
| *Hs*RPA-eGFP | RPA70,<br>RPA32,<br>RPA14 |  | *E. coli* | 8xHis-<br>eGFP-RPA70 | O. Belan et al. 2021<sup>48</sup> |

### Optical tweezer DNA substrates

#### Unconstrained biotinylated λ-DNA

- Lambda DNA (0.3 μg/μL, ThermoFisher Scientific)
- Biotin-14-dATP (Invitrogen)
- Biotin-14-dCTP (Invitrogen)
- dGTP (Invitrogen)
- dTTP (Invitrogen)
- Klenow Fragment (3’→ 5’ exo-) (NEB)
- QIAquick PCR Purification Kit (QIAGEN)

#### End-capped biotinylated λ-DNA

- Unmethylated Lambda DNA (Promega)
- End-cap 1 (IDT, HPLC purified) 5′ AGG TCG CCG CCC GGA GTT GAA CG**T T**T**T**T**T** ACG TTC AAC TCC 3′ End-cap 2 (IDT, HPLC purified) 5′ GGG CGG CGA CCT CAA GTT GGA CAA **T**T**T**T**T T**TG TCC AAC TTG 3′

- Bases in bold are biotinylated
- T4 polynucleotide Kinase (NEB)
- T4 DNA Ligase (NEB)
- PreCR Repair Mix (NEB)

### Expression and purification of TRR and TOP3-RMI1_1-219_

Human TRR (full length TOP3α-RMI1-RMI2) and TOP3-RMI1_1-219_ were recombinantly expressed using the biGBac expression system and purified from Sf9 insect cells. Each complex was co-expressed with their respective RMI1/2 variants in a pBIG1a plasmid assembled from individual pLIB plasmids, with a C-terminal ybbR-Twin Strep tag on TOP3α.

Both complexes were purified with protocols modified from ^30,33,49,50^. Sf9 cells were harvested after 3 days of infection, washed with PBS, and frozen at -80°C until purification. The cells were thawed in 30 mL strep wash buffer (50 mM Tris-HCl pH 7.5, 200 mM NaCl, 5 mM MgCl_2_, 10% (v/v) glycerol, 1 mM DTT) per 5 mL pellet supplemented with 1 protease inhibitor tablet (cOmplete Protease Inhibitor Cocktail EDTA-free tablets, Roche) per 30 mL of lysate, 300 mM NaCl, and Salt Active SuperNuclease (SinoBiological). The resuspended cells were dounce homogenised and sonicated before centrifugation at 40,000 × g for 45 minutes. The supernatant was filtered through a 0.45 μm syringe filter before loading on a 5 mL StrepTrap XT column (Cytiva) equilibrated with wash buffer and eluted with 50 mM biotin. The eluted fractions were pooled and diluted 1:1 with heparin wash buffer (50 mM Tris-HCl pH 7.5, 5% (v/v) glycerol, 1 mM DTT) and then loaded on a 1 mL HiTrap Heparin HP (Cytiva) column equilibrated with 10% heparin elution buffer (heparin wash buffer plus 1 M NaCl). The complex was eluted with a linear gradient of 100 mM – 1 M NaCl. For TRR, the fraction with the highest concentration of protein was used for Sfp-mediated ybbR labelling with AlexaFluor 555 (A555) for 2 hours at room temperature in gel filtration buffer (50 mM Tris-HCl pH 7.5, 200 mM NaCl, 10% (v/v) glycerol, 0.1 mM EDTA, 1 mM DTT) supplemented with 10 mM MgCl2. For TOP3-RMI1_1-219_, all complex containing fractions were pooled and concentrated before labelling with Cy3B overnight at 4°C. The labelling reaction was loaded onto a 10/300 GL Superdex 200 column (Cytiva) for gel filtration. The concentration and labelling efficiency of the complex containing fractions were measured via nanodrop and the highest concentration fraction was aliquoted in small volumes and flash frozen in liquid nitrogen for storage at -80°C.

### Expression and purification of RPA-eGFP

Human RPA-eGFP hetero-trimeric complex was expressed in *E. coli* from a plasmid kindly donated by Simon Boulton^48^, which includes an N-terminal eGFP and 8x-His-tag on RPA70. Expression was induced at an OD_600_ of 0.5 with 0.4 mM IPTG at 18°C for 16 hours. The cells were harvested at 3,600 × g and washed with 3X pellet volumes of PBS before storing at -80°C. The pellet was resuspended in 5X pellet volume of lysis buffer (20 mM Tris-HCl pH 7.5, 500 mM NaCl, 2 mM MgCl_2_, 10% (v/v) glycerol, 2 mM β-mercaptoethanol, 50 mM imidazole) supplemented with two protease inhibitor tablets (Roche) and 300 units of Benzonase (Novagen). The cells subjected to sonication before centrifugation at 40,000 × g for 45 minutes. The supernatant was syringe filtered (0.45 μm) and loaded on a 5 mL HisTrap column (Cytiva) equilibrated with wash buffer (20 mM Tris-HCl pH 7.5, 400 mM NaCl, 10% (v/v) glycerol, 2 mM β-mercaptoethanol, 10 mM imidazole). The column was washed with 5 column volumes of wash buffer and the protein eluted with elution buffer (20 mM Tris-HCl pH 7.5, 250 mM NaCl, 10% (v/v) glycerol, 2 mM β-mercaptoethanol, 250 mM imidazole). hRPA-eGFP containing fractions were pooled and diluted 1:4 in heparin wash buffer (20 mM Tris-HCl pH 7.5, 50 mM NaCl, 10% (v/v) glycerol, 2 mM β-mercaptoethanol). The diluted sample was loaded onto a 5 mL HiTrap Heparin HP column (Cytiva) equilibrated with heparin wash buffer and eluted with a linear gradient of 10-50% elution buffer (20 mM Tris-HCl pH 7.5, 1 M NaCl, 10% (v/v) glycerol, 2 mM β-mercaptoethanol). RPA-eGFP containing fractions were pooled, concentrated, quantified, and flash frozen in liquid nitrogen for storage at -80°C.

### TOP3α relaxation assay

200 nM of TRR or TOP3-RMI1_1-219_ were incubated with 0.5 ng of negatively supercoiled pBR322 (Inspiralis) plasmid at 37°C for 1 hour in 20 mM Tris-HCl pH 7.5, 125 mM potassium glutamate, 2.5 mM MgCl_2_, and 1 mM DTT. The reaction was stopped with a final concentration of 0.5% (v/v) SDS and 1.6 U proteinase K for 30 minutes at 37°C before loading onto a 1% 1x TBE agarose gel free of intercalating dye. The samples were run alongside 0.5 ng of relaxed pBR322 and protein free negatively supercoiled pBR322 at 80V for 2 hours, then stained with SYBR safe dye for 15 minutes before imaging.

### Preparation of unconstrained λ-DNA

Biotinylated λ-DNA with open ends were generated by filling in the overhangs with the 3’-5’ exonuclease Klenow fragment with GTP, TTP, Biotin-14-ATP and Biotin-14-CTP. Following the reaction at 37°C and enzyme deactivation at 70°C, the biotinylated DNA was purified using the QIAquick PCR purification kit according to the vendor-supplied protocol.

### Preparation of end-capped λ-DNA

Biotinylated end-capped λ-DNA was generated as previously described^47^. Briefly, the biotinylated end-caps were phosphorylated using polynucleotide kinase. The phosphorylated end-caps were then ligated to λ-DNA using T4 DNA ligase. 25 μL of the end-capped product was used with the PreCR repair kit to ensure a majority of the DNA is free of nicks. 5-10 μL of this reaction was then used to generate the negatively supercoiled DNA in the C-trap via the ODS method.

### Negative supercoiling generation by ODS

To enable ODS, a DNA molecule with end-closed (looped) ends is constructed by ligating ‘end-caps’, containing multiple biotin moieties, to each end of the DNA. The end-capped molecule is held in a torsionally constrained state when at least two biotins on each end-cap are bound to streptavidin-coated optically trapped beads. ODS exploits the fact that, at high forces (typically > 100 pN), the biotin-streptavidin bonds are less stable. This results in transient breakage of the bonds, such that the DNA molecule is temporarily tethered via a single linkage. Since the DNA is most stable in an underwound state at these high forces, transient disruption of the biotin-streptavidin bonds induces a fixed reduction in the linking number (corresponding to unwinding of the molecule). Subsequent lowering of the force stabilizes the biotin-streptavidin bonds and preserves the molecule in a negatively supercoiled state. The amount of unwinding (negative supercoiling) can vary between 0 > σ > -0.7 and depends on the number of stretch-retract cycles performed and the duration that the molecule is held at high force. The supercoiling density can be determined based on the distance at 70 pN (Fig. S1A), where the absolute uncertainty in the magnitude of σ is estimated as ∼0.03 ^28,47^. However, the uncertainty in determining σ = 0 is smaller (typically < 0.01), owing to the fact that the initial stretch cycle will always begin in a non-supercoiled state. For this reason, we use σ = -0.01 as the cut-off for distinguishing supercoiled and non-supercoiled molecules.

### Single-molecule experiments

Dual trap and quadruple trap (Q-trap) experiments were conducted on a LUMICKS C-trap. Trapping laser was used at 25-30% power for dual trap experiments and 75-80% power with a split of 67% across traps 1 (beads 1-2) and 2 (beads 4-3) for Q-trap experiments, both achieving a trap stiffness of 0.2-0.3 pN/nm. Images were acquired with a pixel size of 0.1 μm x 0.1 μm and a pixel dwell time of 0.00012 seconds. The 2-D scan frame rate was ∼5.7 seconds. Fluorescent lasers (532 nm and 488 nm) were used at ∼ 1 μW power. ∼4.4 μm diameter streptavidin coated beads (Spherotech) were used for all experiments.

All experiments were conducted within a 5-channel microfluidics chip which was passivated with 0.5% Pluronic F-127 solution. Channels 1-3 contained 20 mM Tris-HCl pH 7.5 and 50 mM NaCl. Channels 4-5 were additionally passivated with 2 mg/mL BSA in protein buffer (50 mM Tris-HCl pH 7.5, 125 mM potassium glutamate, 2.5 mM MgCl_2_) for 45 minutes before proteins were introduced in 0.5 mg/mL BSA in protein buffer plus 1 mM DTT. RPA-eGFP was used at 5 nM, while TRR-A555 and TOP3α-RMI1_1-219_ were used at 2 nM.

Right-handed braided substrates were generated as described in Cutts, E. *et al.* 2025^22^. For all braiding experiments, a manual FE curve for both DNAs was collected in channel 3 before and after braiding, and after incubation with proteins in either channel 4 or channel 5. For RPA binding experiments, braids were transferred to the channel containing RPA-eGFP with both bead pairs at 10 μm in the x-axis for a force of 0 pN in trap 1 (bead 1) and trap 2 (bead 4). Bead pair 1-2 and bead pair 4-3, respectively, were arranged at the top and bottom of a ∼18 μm long 2-D confocal scan ROI. Once in the RPA containing channel, beads 2 and 3 were individually moved rightward in the x-axis to generate force on beads 1 and 4 respectively. Each bead was moved in alternating increments to maintain a similar force in both traps. After RPA binding was observed at the braid centre, the braid was moved to a channel containing TRR or Top3α-RMI1_1-219_ for decatenation experiments.

Mean resolution probability was calculated as previously described^22^ for braid resolutions occurring withing a 2-minute sampling time. Decatenations occurring within the sampling time were considered a success (k) out of a number of trials (n), and α = k + 1 and β = n – k +1. The mean resolution probability was calculated as mean 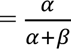 and 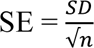, where SD = 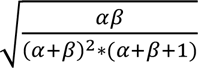.

Kymographs of 2D-scans were generated from a 1-pixel wide segmented line ROI between beads 1 and 3, using the RPA-eGFP signal as a guide for the DNA location. The Dynamic Kymograph Fiji plugin (https://github.com/rudyzhou/Dynamic_Kymograph) was used to stitch the pixels from each frame of the 2D-scan together into a kymograph. The resulting kymograph can be correlated to the plot of force over time to determine the force at which RPA binding is first detected near the centre of the braid. The Fiji ImageJ plot profile function was used to extract the normalised pixel intensity in the blue and green channels for specific frames of the kymograph to demonstrate that the fluorescence signal corresponds to the centre of the braid.

## Acknowledgments

We thank members of the DNA Motors and Single Molecule Imaging groups (MRC LMS), especially Gemma Fisher, Korak Ray, and Paul Girvan, for critical reading of the manuscript. This work was funded by the Medical Research Council (UKRI MC-A652-5PY00).

## Author contributions

L.A. and S.S. conceived the study. A.I.K. purified RPA. S.S. purified the proteins, performed the biochemical analysis of protein samples, performed all single-molecule experiments, and performed all data analysis. G.A.K. provided protocols and critical advice for supercoiling assays. D. R. provided advice for DNA gap assays. L.A. wrote the manuscript. L.A. led the research and S.S. performed analysis and interpretation of the results. G.A.K., A.I.K and S.S. discussed the results, commented and edited the manuscript.

## Inclusion and Ethics

This study did not involve human participants, human data, or animal subjects. All experimental procedures were conducted in compliance with institutional and national regulations. No ethical issues related to inclusion or research integrity were identified.

## Data and material availability

All reagents will be available upon reasonable request. All raw data and custom scripts for single-molecule data analysis will be available on Zenodo upon publication.

## Supplementary figures

**Figure S1.**
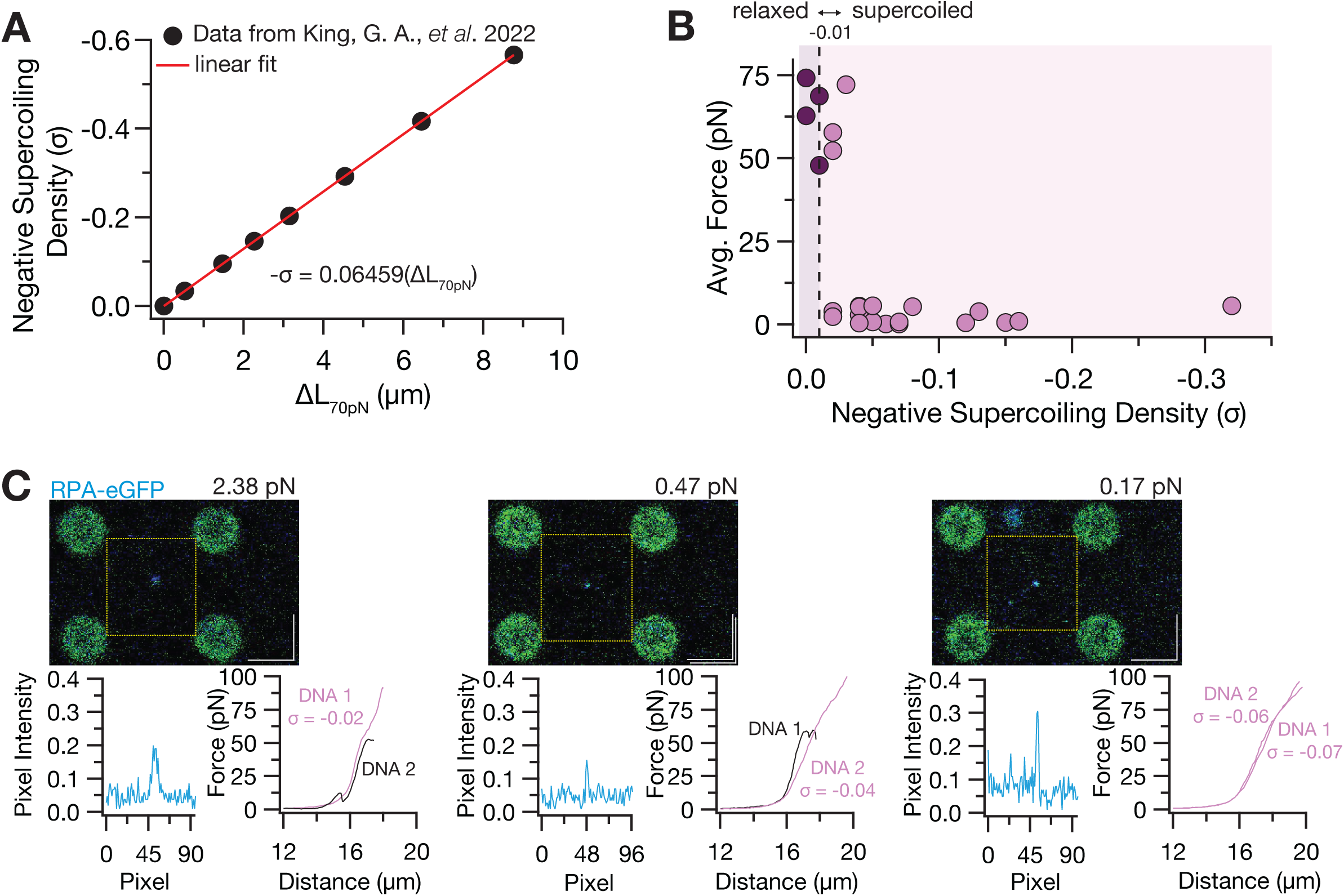
Negative supercoiling in DNA braids lowers the force threshold for RPA binding. **(A)** The linear function used to calculate the supercoiling density of the constrained DNAs used in this study. Original data for σ for a given difference in DNA extension at 70 pN between relaxed and negatively supercoiled DNA from King, G.A. *et al.* 2022 ^46^. **(B)** Correlation of the average force in trap 1 and 2 at which RPA bound to the braid to the supercoiling density of the DNA in the braid. The dotted line at σ = -0.01 denotes the cutoff for relaxed DNA. **(C)** More examples of RPA binding at the braid at low force in braids composed of one or both negatively supercoiled DNAs. A representative frame from different 2-D scans is shown with the corresponding average force in Trap 1 and 2. The blue pixel intensity is shown for the ROI denoted by the dotted white box. FE curves show the type of DNA between each bead pair and the calculated supercoiling density (σ) if the DNA is a constrained molecule. Black denotes a nicked DNA and light purple for negatively supercoiled DNA.

**Figure S2.**
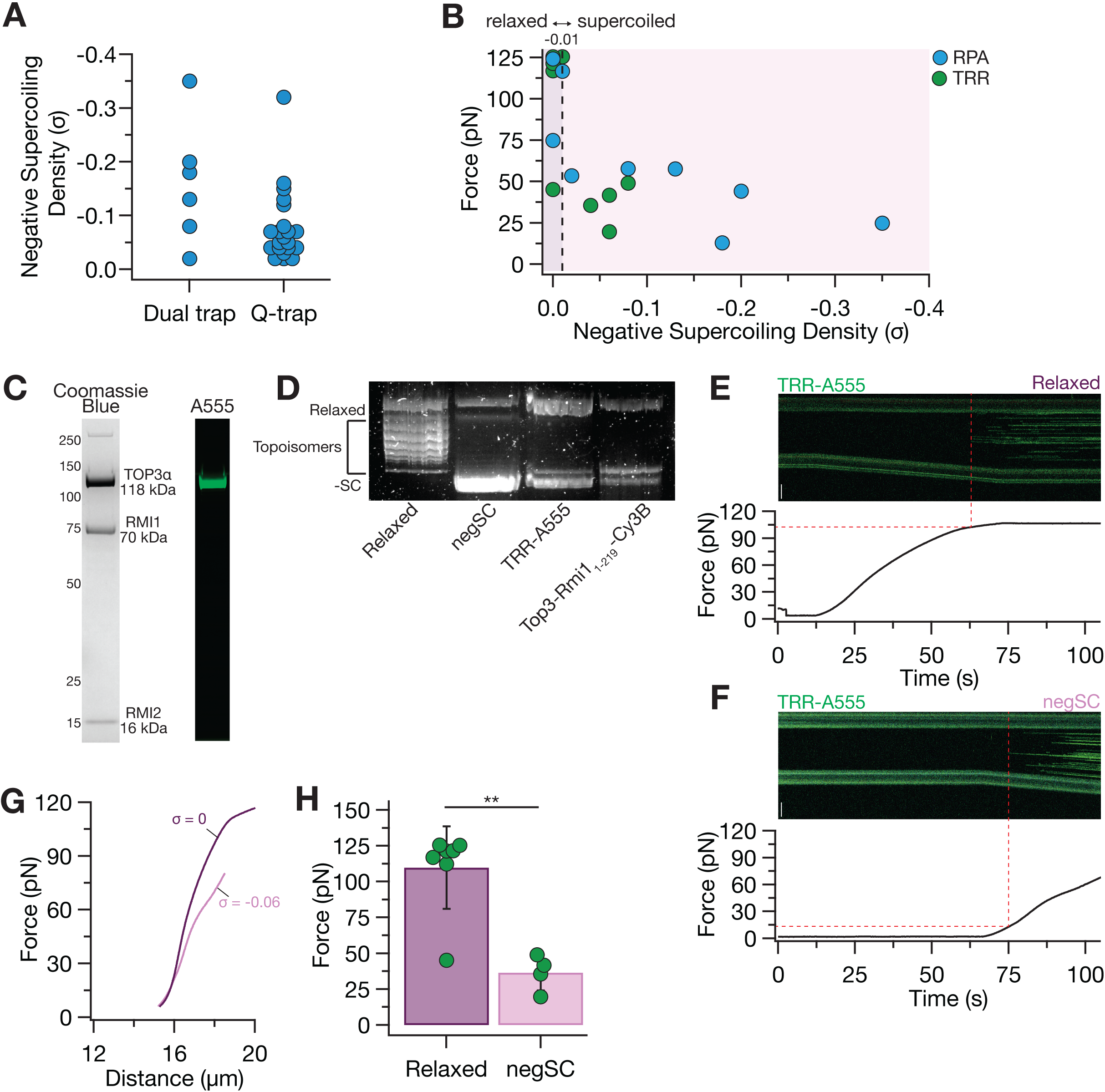
RPA and TRR binding to negatively supercoiled linear DNA. **(A)** Distribution of the supercoiling densities of the DNAs used in the dual trap vs the Q-trap experiments in this study. **(B)** Correlation of the force on bead 2 at which RPA (blue) or TRR (green) bound to a single DNA to the supercoiling density of the DNA in dual-trap experiments. The dotted line at σ = -0.01 denotes the cutoff for relaxed DNA. **(C)** SDS-PAGE of TRR-A555 after 3-step purification. (left) shows the Coomassie blue staining and (right) shows the corresponding green channel fluorescence signal of the A555 labelled TOP3α. **(D)** Agarose gel image of negatively supercoiled plasmid relaxation assay. The first two lanes show protein-free controls of relaxed and negatively supercoiled pBR322 plasmid. The last two lanes show the relaxation activity of the purified TRR-A555 and TOP3-RMI1_1-219_ by a depletion of the negatively supercoiled substrate and an increase in the relaxed substrate. **(E-F)** Example kymograph of a constrained, relaxed DNA (F) and negatively supercoiled DNA (σ = -0.06) (G) extended in a channel containing 2 nM TRR-A555 with corresponding force. Dotted red lines indicate the force at which TRR bound. Scale bars = 5 μm. **(G)** FE curves of the DNAs shown in panels F and G taken in the TRR-A555 channel during the kymographs. Since TRR can relax the negatively supercoiled DNA at low force, the supercoiling density was measured by the extension of the DNA at 70 pN in the protein channel. **(H)** Comparison of the force at which TRR bound to a single relaxed or supercoiled DNA. Mean force in trap 2 reported with SD (error bars): Relaxed = 109.74 pN ± 28.85 pN (n = 7), negSC = 36.47 pN ± 12.47 pN (n = 4). ** denotes a two-tailed p-value of 0.0011 as calculated by an un-paired *t* test.

**Figure S3.**
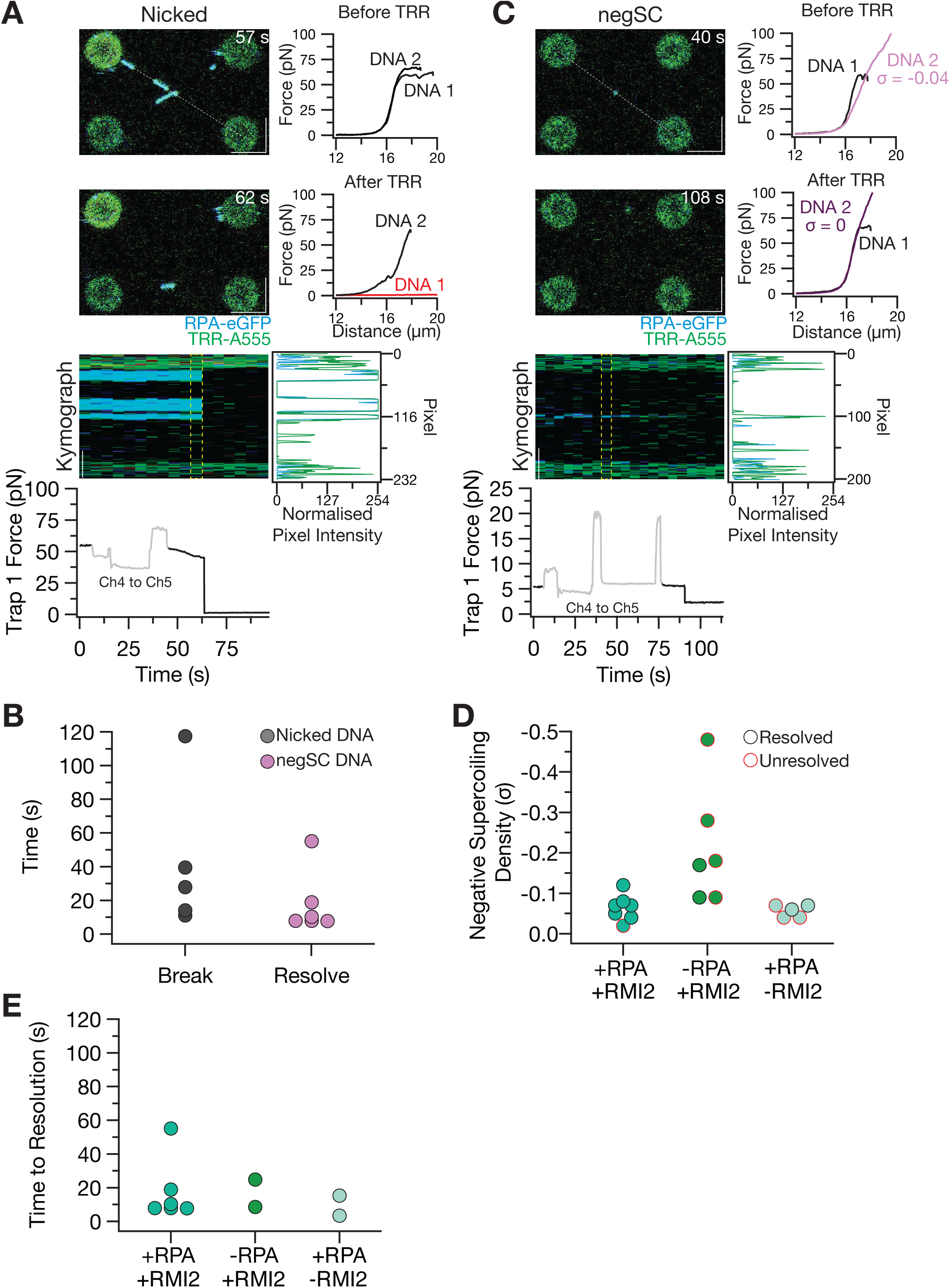
TRR decatenation depends on substrate and presence of RPA and RMI2. **(A)** Additional example of a braid formed with two nicked DNAs. **(B)** Time required for breakage or resolution within the sampling time of 2 minutes in the channel containing 2 nM TRR. **(C)** Additional example of a braid formed with one negatively supercoiled DNA (σ = -0.04). **(D)** The supercoiling densities of the negatively supercoiled DNA in the braids exposed to TRR ± RPA or ± RMI2. Red outline indicates the braid was unresolved at 5 pN within the 2-minute sampling time, black outline indicates the braid was resolved at 5 pN within the 2-minute sampling time. **(E)** Time in seconds from entry into the channel containing TRR ± RPA or ± RMI2 to resolution at 5 pN for the braids that were resolved within the 2-minute sampling time.

